# Systems Biology behind immunoprotection of both Sheep and Goats after Sungri/96 PPRV vaccination

**DOI:** 10.1101/2020.08.14.252056

**Authors:** Sajad Ahmad Wani, Manas Ranjan Praharaj, Amit R Sahu, Raja Ishaq Nabi Khan, Kaushal Kishor Rajak, Dhanavelu Muthuchelvan, Aditya Sahoo, Bina Mishra, R. K. Singh, Bishnu Prasad Mishra, Ravi Kumar Gandham

## Abstract

Immune response is a highly coordinated cascade involving all the subsets of PBMCs. In this study, RNA-Seq analysis of PBMC subsets - CD4+, CD8+, CD14+, CD21+ and CD335+ cells from day 0 and day 5 of Sungri/96 Peste des Petits Ruminants vaccinated sheep and goats was done to delineate the systems biology behind immune - protection of the vaccine in sheep and goats. Assessment of the immune response processes enriched by the differentially expressed genes in all the subsets suggested a strong dysregulation towards development of early inflammatory microenvironment, which is very much required for differentiation of monocytes to macrophages, and for activation and migration of dendritic cells into the draining lymph nodes. The protein - protein interaction networks among the antiviral molecules (IFIT3, ISG15, MX1, MX2, RSAD2, ISG20, IFIT5 and IFIT1) and common DEGs across PBMCs subsets in both the species identified ISG15 to be an ubiquitous hub, that helps in orchestrating antiviral host response against PPRV. IRF7 was found to be the key master regulator activated in most of the subsets in sheep and goats. Most of the pathways were found to be inactivated in B - lymphocytes of both the species indicating that 5 dpv is too early a time point for the B - lymphocytes to react. The cell mediated immune response and humoral immune response pathways were found more enriched in goats than in sheep. Though, animals from both the species survived the challenge, a contrast in pathway activation was observed in CD335+ cells.

**Importance:** Peste des petits ruminants (PPR) by PPRV is an OIE listed acute, contagious transboundary viral disease of small ruminants. Attenuated Sungri/96 PPRV vaccine used all over India against this PPR, provides long-lasting robust innate and adaptive immune response. The early antiviral response was found mediated through type I interferon independent ISGs expression. However, systems biology behind this immune response is unknown. In this study, in vivo transcriptome profiling of PBMC subsets (CD4+, CD8+, CD14+, CD21+ and CD335+) in vaccinated goats and sheep (at 5 days of post vaccination) was done to understand this systems biology. Though there are a few differences in the systems biology across cells (specially the NK cells) between sheep and goats, the co-ordinated response that is inclusive of all the cell subsets was found to be towards induction of strong innate immune response, which is needed for an appropriate adaptive immune response.

## 1. Introduction

Peste des petits ruminants (PPR) is an OIE listed acute, highly contagious transboundary viral disease of small ruminants, caused by PPR virus of genus *Morbillivirus* and family *Paramyxoviridae* [1]. Morbidity and mortality can be as high as 100% and 90%, respectively [2]. The disease manifests as fever, discharge from the eyes and nose, stomatitis, pneumonia and enteritis [3]. PPRV vaccine developed by continuous passage (N=59) of Sungri/96 strain in Vero cells is widely used throughout India [4]. The vaccine provides long-lasting robust innate and adaptive - humoral and a strong cell-mediated immunity [2],which, however, warrants further investigation [5], [6], [7]. PPRV is lymphotropic and epitheliotropic [8], [9], [10]. The primary receptors for PPRV include the signalling lymphocyte activation molecule (SLAM) on activated T cells, B cells, and dendritic cells and Nectin-4 receptor on epithelial cells [8], [9], [11].

Immune response is complex within a host and different cell types respond differently to infection as different classes of receptors receive cues and produce distinct effector molecules [12]. It is a highly coordinated effort of distinctly programmed hematopoietic cell types, and a product of various direct and indirect effects and interactions between similar or different cell types [13]. Moreover, the tissue microenvironment also affects the elicited immune response. In case of viruses, the complexity of the host response depends on variations in genetic makeup, cell tropism and replication kinetics [12], [13], [14], [15]. PBMCs include T helper cells (CD4+), T cytotoxic cells (CD8+), B-lymphocytes (CD21+), monocytes(CD21+), natural killer cells(CD335+) and dendritic cells (CD320+), which play an important role in virus recognition and induce immune response for host defence. While analysing whole blood or PBMCs, the response of under-represented cell populations can be masked [13].

Despite advances in our understanding of vaccines, the mechanisms by which protective immune responses are orchestrated among the cell subsets are little known. Molecular patterns and gene signatures detected in blood post vaccination represent a strategy to prospectively determine vaccine efficacy [16]. The conventional immunological methods like ELISA, ELISpot etc. are of utmost importance in this regard and may continue to remain so in future [17]. However, these approaches are inept at predicting the systems biology behind immune - protection. Delineating the systems biology would help understanding the molecular mechanisms of vaccine induced immune responses. RNA sequencing is a widely used quantitative transcriptome profiling system for deciphering the systems biology comprehensively [18]. Previously, RNA sequencing was used to unravel transcription factors, which modulate immune response to PPRV Sungri/96 live attenuated vaccine strain *in vitro* in PBMCs [6]. Also, a predicted immune signalling pathway of PPRV Sungri/96 vaccine induced immune response with predominant role of IRFs, TRIMs and ISGs in creation of robust antiviral state *in vitro* in PBMCs has been proposed [7].

Till now there are no *in vivo* reports of transcriptome profiling of PBMC subsets in PPRV vaccinated goats and sheep. Herein, transcriptional profiling of circulating CD4+, CD8+, CD14+, CD21+ and CD335+ cells of PPRV vaccinated sheep and goats at 0 day (control i.e. just before vaccination) and before the development of antibody response (5 days post vaccination i.e. 5dpv) to decipher the vaccine induced immune response was done.

## 2. Materials and methods

### 2.1 Animal experiment, ethics statement and virus

Live attenuated PPR vaccine virus (Sungri/96) was used as vaccine virus. Permission for studies on animal subjects was obtained and protocols approved from IVRI – Institutional Animal Ethics Committee (IAEC) under CPCSEA, India vide letter no. 387/CPCSEA. The vaccine potency testing experiment was carried out as per the guidelines of Indian Pharmacopeia – 2014 (page.no: 3626).

In this study, healthy sheep and goats (n=5, age=12 months) confirmed negative for PPRV antibodies (c-ELISA and SNT) and PPRV antigen (s-ELISA) [19], [20] were used. On c-ELISA, the samples with a percent inhibition (PI) value of >40% were considered positive. The animals were acclimatised for 14 days, followed by vaccination on day 0 with a 10^3^ TCID50 field dose of Sungri/96 strain through sub-cutaneous route, as mentioned in our previous report [21]. All the animals vaccinated survived the challenge from the virulent PPRV in the vaccine potency testing experiment.

### 2.2 Isolation of T helper cells, T cytotoxic cells, B lymphocytes, monocytes and natural killer cells by magnetic assisted cell sorting technology (MACS)

Blood was collected from the animals (n=5) in heparin coated vacutainer vials at 0 day (just before vaccination) and 5 days post vaccination (5 dpv). PBMCs were isolated by using Ficol Histopaque gradient method. PBMCs were strained through cell strainer of 0.40-micron size. The PBMCs cell subsets were enriched by positive selection using indirect magnetic assisted cell sorting technology (Miltenyi Biotech). Cell sorting was done as per the manufacturers protocol. Initially, the cell-specific surface marker FITC-conjugated primary antibodies, anti CD4+ (T helper cells, #MCA2213F), anti CD8+ (T cytotoxic cells, #MCA2216F), anti CD14+ (Monocytes, #MCA1568F), and anti CD21+ (B lymphocytes, #MCA1195F), were used. For CD335 (NK cell) anti CD335+ (#MCA5933GA as primary antibody) and FITC labelled secondary antibody (#F9137) were used. Subsequently, the cells were magnetically labeled with anti - FITC MicroBeads. Then the cell suspension was loaded on a miniMACS^®^ column which was placed in the magnetic field of a MACS Separator. The magnetically labeled cells were retained in the column while the unlabeled cells run through. After removal of the column from the magnetic field, the magnetically retained cells were eluted as positively selected cell fraction. The purity of the cells was further checked by flow cytometer. The cells were stored in RNA later for further use at −80°C. Cells were kept on ice and cold buffers were employed to minimize alterations in gene expression during labelling and sorting.

### 2.4 RNA-sequencing of the samples

Total RNA from each of the PBMC subsets was isolated using the RNeasy Mini kit (Qiagen GmbH, Germany) as per the manufacturer’s protocol. The integrity and quantity of isolated RNA were assessed on a Bioanalyzer (Agilent Technologies, Inc). The library was prepared using NEBNext Ultra RNA Library Prep Kit for Illumina (NewEngland Biolabs Inc.) following the manufacturer’s protocol. Approximately, 100ng of RNA from each sample was used for RNA library preparation. The quality of the libraries was assessed on Bioanalyzer. Libraries were quantified using a Qubit 2.0 Fluorometer (Life technologies) and by qPCR. Library (1.3ml, 1.8pM) was denatured, diluted and loaded onto a flow cell for sequencing. cDNA library preparation and Illumina Sequencing was performed at Bioserve Pvt. (Hyderabad, India). RNA-Seq data was generated in FASTQ format.

### 2.5 Raw data processing

Raw sequence data from each sample was subjected to quality control checks using FastQC (Babraham Bioinformatics). Moreover, low quality reads with a mean phred score less than or equal to 25 and reads shorter in length than 50 nt were removed using prinseq-lite software [22] before downstream analysis. The data was submitted to the GEO database with accession number GSE155504.

### 2.6 Differential expression and identification of differentially expressed genes (DEGs)

Figure 1 summarizes the steps used in the analysis. Quality filtered reads from control and vaccinated samples (0 day and 5 dpv) were mapped to the *Capra hircus* or *Ovis aries* reference genome for the respective subsets. The gene counts were obtained using Bowtie2.0 in RSEM [23]. The counts were used for calculating differentially expressed genes (DEGs) by use of R packages - EBSeq, DESeq2, edgeR. The common DEGs from the three packages were used for downstream analysis while fold changes for the corresponding genes were taken from DESeq2 [24].

**Figure 1:**
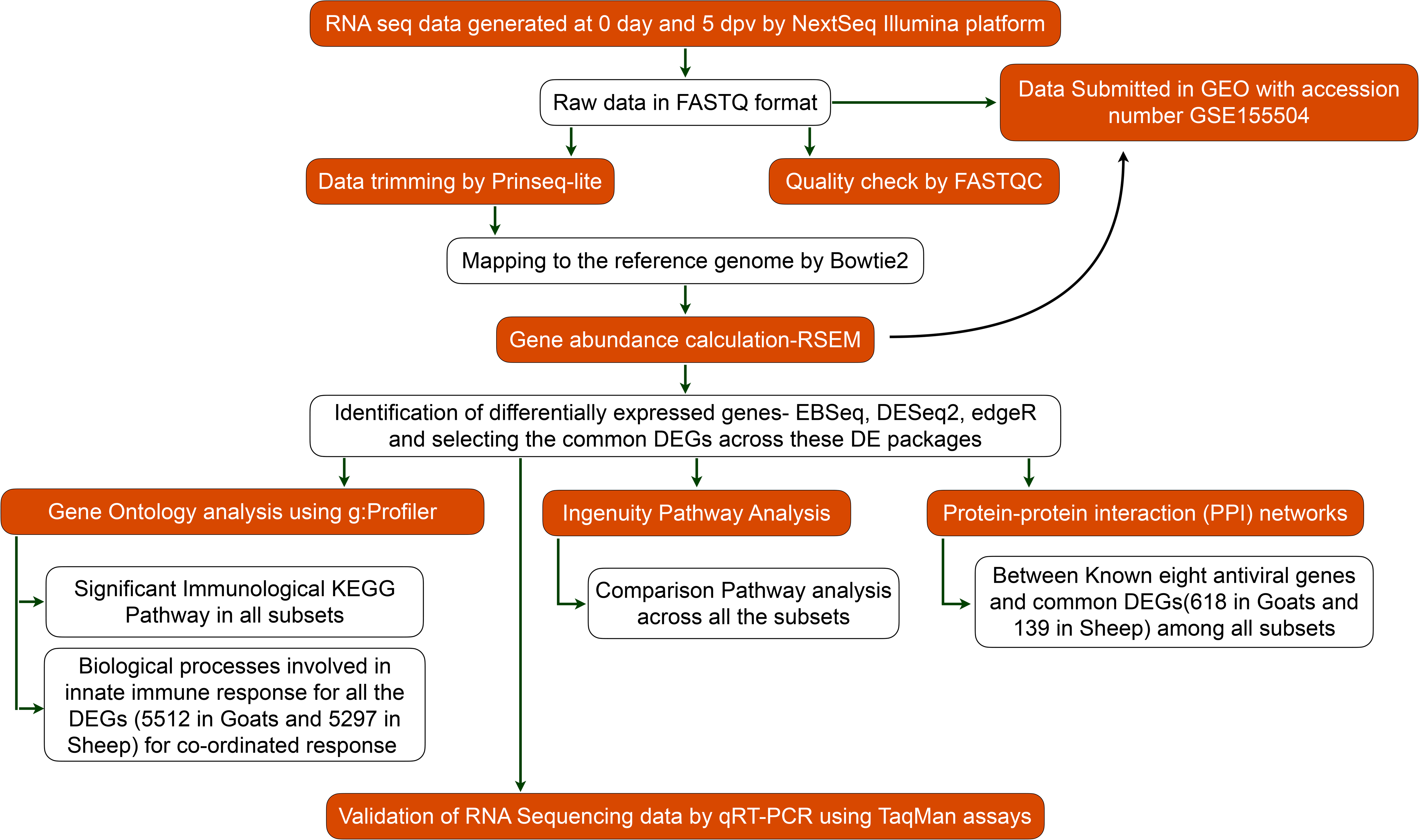
Workflow for RNA sequencing data analysis

### 2.7 Gene Ontology Analysis

Initially, DEGs of each subset (CD4+, CD8+, CD14+, CD21+ and CD335+) in sheep and goats were functionally annotated in g:Profiler to identify the significant immune system KEGG pathways. The expression of common DEGs of each subset between sheep and goats that are involved in immunological KEGG pathways is represented in a heatmap. Finally, to understand the co-ordinated response across all the subsets, genes expressed in all cell subsets were functionally annotated in g:Profiler (a gene is considered expressed if it is expressed in one subset).

### 2.8 Comparison analysis using Ingenuity Pathway Analysis (IPA) analysis

Ingenuity Pathway Analysis (IPA) IPA is an all-in-one, web-based software application that enables analysis, integration, and understanding of data from gene expression, miRNA, and SNP microarrays, as well as metabolomics, proteomics, and RNAseq experiments.The DEGs from all the subsets in both species were overlaid in IPA against its Ingenuity Knowledge Base (IKB) to perform a comparison analysis. Canonical pathways activated (Z score > 2) or inactivated (Z score < −2) across all the subsets were identified.

### 2.9 Protein-protein interaction networks

Using knowledge based approach [25], [26], antiviral genes - IFIT3, ISG15, MX1, MX2, RSAD2, ISG20, IFIT5 and IFIT1 were selected based on their expression in at least one subset. Protein - protein interactions between these antiviral molecules and the common genes among all the subsets for each species were extracted using STRING [27] and customized scripts. The degree or connectivity was calculated using igraph package [28]. The complete interaction networks were visualized in Cytoscape 3.8.0 [29].

### 2.10 Validation of DE genes by quantitative real time PCR (qRT-PCR)

qRT-PCR was performed using Applied Biosystems 7500 Fast system to validate the expression of key genes using GAPDH as an endogenous control by TaqMan chemistry in PBMC subsets. GAPDH was employed as the internal control as it was found to be suitable endogenous control in earlier studies in PPR [30]. Key genes used in the study for validation by q-RT-PCR and their TaqMan probe IDs are - DDX58, Ch04684385_m1; IFIT3, AIAA1E0; IRF7, AI89L87; MX1, Oa04659431_m1; ISG15, AI70N2Z; and GAPDH, AIFAT31. All the samples were run in triplicates. The relative expression of each sample was calculated using the 2^-ΔΔCT^ method with control as calibrator [31].

## Results

In the present study, CD4^+^, CD8^+^, CD14^+^, CD21^+^ and CD335^+^ cells were enriched **(Supplementary Figure 1)** from the blood collected (5 goats and 5 sheep) at 0 day and 5 dpv (5 days post vaccination). RNA was isolated from these subsets to profile the transcriptome with an aim to delineate the systems biology behind the Sungri/96 vaccine induced immuno-protection at 5 dpv in sheep and goats. The number of DEGs in CD4^+^, CD8^+^, CD14^+^, CD21^+^ and CD335^+^ cells were 1834, 1641, 2343, 3910 and 3607, respectively, in goats and 1464, 1586, 1847, 721 and 4019, respectively, in sheep **(Figure 2A and 2B)**. Venn diagram was generated to examine the common and unique DEGs among cells. On comparison, 618 and 139 DEGs were found common among CD4^+^, CD8^+^, CD14^+^, CD21^+^ and CD335^+^ cells, in goats and sheep, respectively. The number of unique DEGs was highest in CD21^+^ cells of goats and CD335^+^ cells of sheep **(Figure 2C and 2D)**.

**Figure 2:**
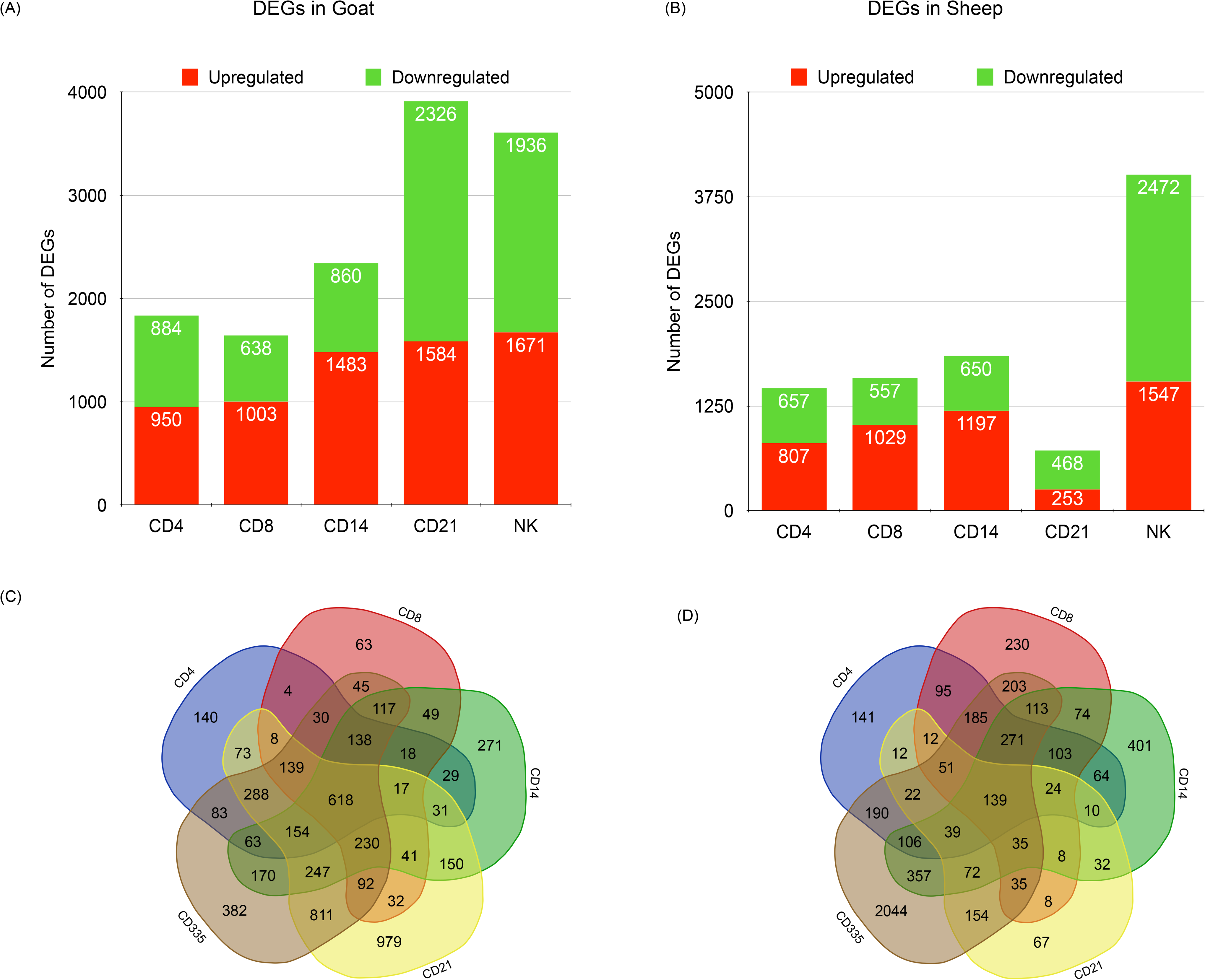
Number of dysregulated DEGs in PBMC subsets (A) Goats and (B) Sheep. Green colour represents downregulation and red colour represents upregulation. (C)Venn diagrams representing unique/common DEGs among cells in goats (D) Venn diagrams representing unique/common DEGs among cells in sheep

### Gene Ontology Analysis

Initially to evaluate the changes within a subset, functional annotation for genes expressed in each subset was done using g:profiler. The immune system KEGG pathways enriched in each subset were assessed. In all the cells an innate immune response leading to cell mediated adaptive immune response was observed **(Figure 3 and Figure 4)**.

**Figure 3:**
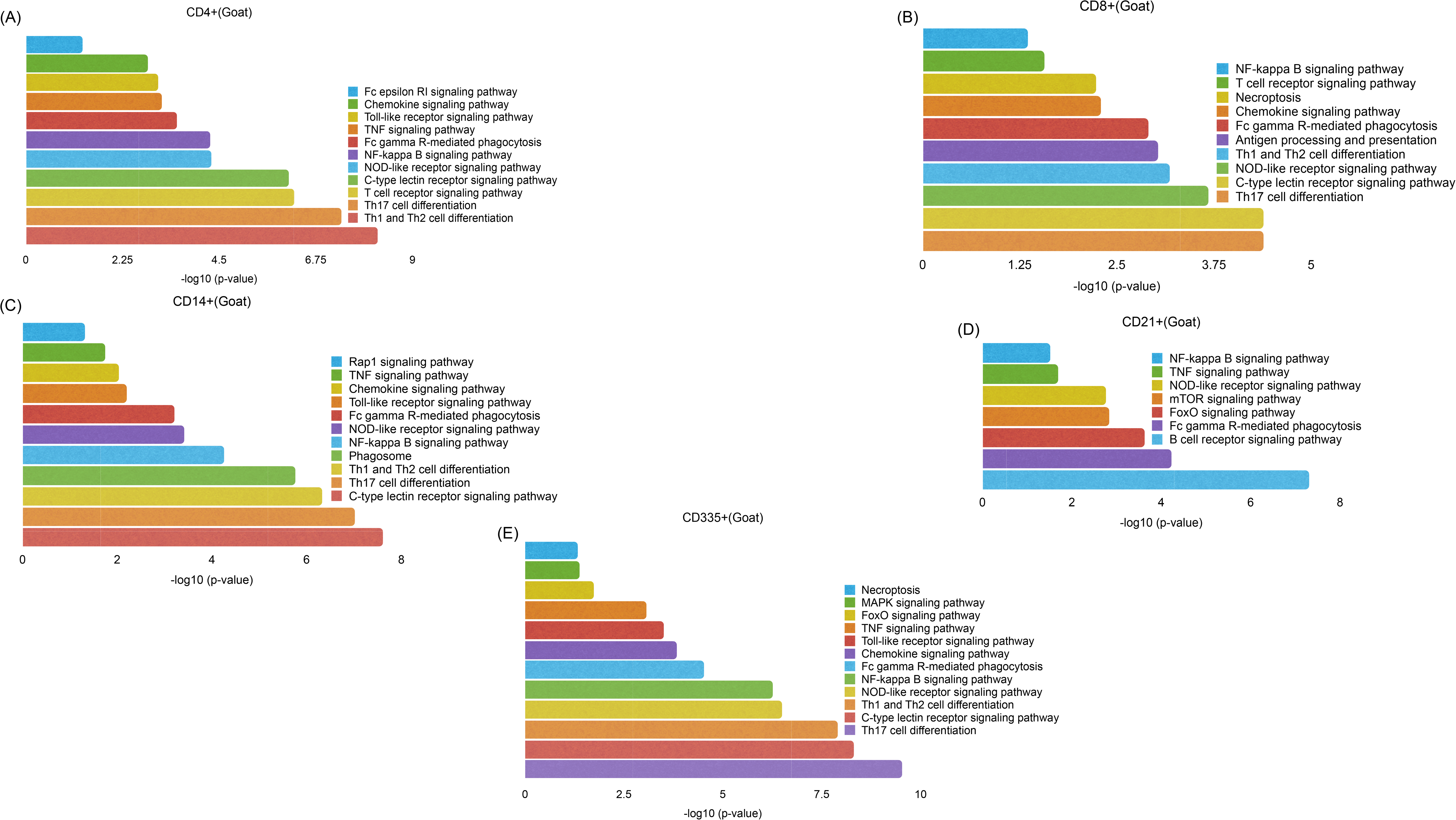
Functional annotation of DEGs involved in immunological processes for each subset of PBMCs: (A) CD4+, (B) CD8+, (C) CD14+, (D) CD21+ and (E) CD335+ using g:Profiler in Goats

**Figure 4:**
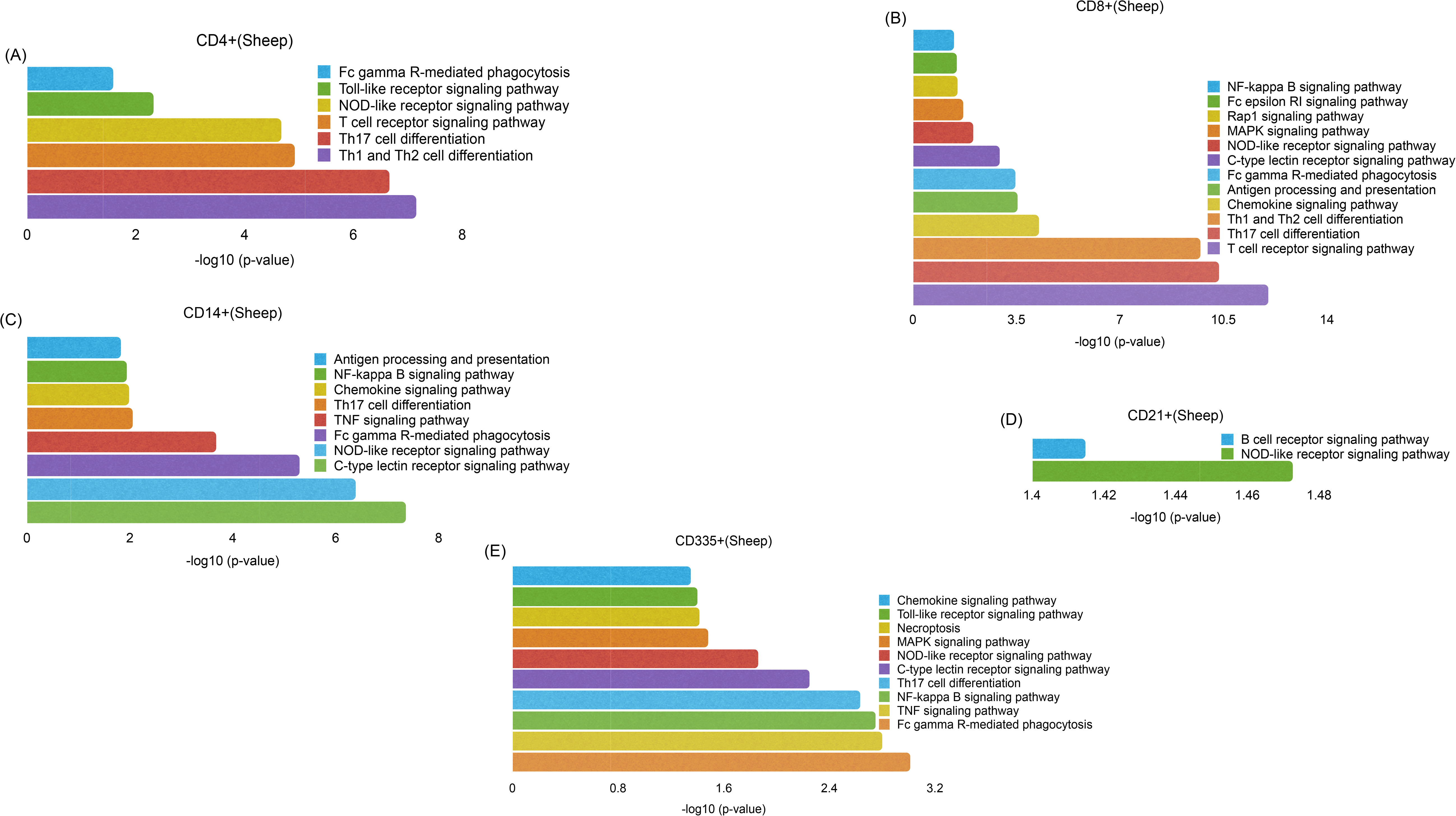
Functional annotation of DEGs involved in immunological processes for each subset of PBMCs: (A) CD4+, (B) CD8+, (C) CD14+, (D) CD21+ and (E) CD335+ using g:Profiler in Sheep

#### CD4+ cells of Sheep and Goats

On comparing CD4+ cells in sheep and goats, Fc gamma R-mediated phagocytosis, Toll-like receptor signaling pathway, NOD-like receptor signaling pathway, Th1 and Th2 cell differentiation, T cell receptor signaling pathway and Th17 cell differentiation were found significantly enriched in both the species. Besides these, in goats, Fc epsilon RI signaling pathway, C-type lectin receptor signaling pathway, TNF signaling pathway, Chemokine signaling pathway and NF-kappa B signaling pathway were found enriched in CD4+ cells.

#### CD8+ cells of Sheep and Goats

In CD8+ cells,Th17 cell differentiation, C-type lectin receptor signaling pathway, NOD-like receptor signaling pathway, Antigen processing and presentation,Th1 and Th2 cell differentiation, T cell receptor signaling pathway, Fc gamma R-mediated phagocytosis, Chemokine signaling pathway, NF-kappa B signaling pathway were found enriched in both sheep and goats. Besides these, three more pathways - Rap1 signaling pathway, Fc epsilon RI signaling pathway and MAPK signaling pathway were enriched in sheep.

#### CD14+ cells of Sheep and Goats

In CD14+ cells of both the species, Th17 cell differentiation, C-type lectin receptor signaling pathway, NOD-like receptor signaling pathway, TNF signaling pathway, Fc gamma R-mediated phagocytosis, Chemokine signaling pathway and NF-kappa B signaling pathway were found enriched. Additionally, in goats, Toll-like receptor signaling pathway, Rap1 signaling pathway and Phagosome Th1 and Th2 cell differentiation were enriched.

#### CD335+ cells of Sheep and Goats

In CD335+ cells, Th17 cell differentiation, Toll-like receptor signaling pathway, C-type lectin receptor signaling pathway, Necroptosis, MAPK signaling pathway, NOD-like receptor signaling pathway, TNF signaling pathway, Fc gamma R-mediated phagocytosis, Chemokine signaling pathway and NF-kappa B signaling pathway were enriched in sheep and goats (**Figure 3 and Figure 4**). Th1 and Th2 cell differentiation and FoxO signaling pathway were found enriched additionally in goats CD335+ cells.

#### Common DEGs in each subset between Sheep and Goats

The common genes in sheep and goats that are involved in immunological processes in each subset are represented in a heatmap (**Supplementary Figure 2**). In CD4+, CD8+, CD14+, CD21+ and CD335+ cells the numbers of common genes involved in immunological processes were found to be 67, 91, 122, 17 and 179, respectively. Most of the common DEGs in CD4+, CD8+ and CD!4+ were found upregulated in both the species. However, in CD21+ and CD335+, a contrast in expression of these genes was observed between sheep and goats. Most of the DEGs were upregulated in goats but downregulated in sheep.

#### Coordinated response

To understand the co-ordinated response across all the subsets, genes expressed in all cell subsets were functionally annotated. A total of 5512 and 5297 genes were found expressed across all subsets **(Supplementary File 2)** in goats and sheep, respectively. Among these, in goats and sheep, 689 and 703 genes, respectively, were found associated with innate immune response biological processes **(Supplementary Figure 3)**. A subset of 544 immune response genes were found to be common between sheep and goats with 144 and 158 genes being unique, respectively. This shows that in both sheep and goats the coordinated vaccine response at 5dpv across all the subsets is towards triggering a strong innate immune response as evident from the upregulation of innate immune genes.

### Comparison analysis across subsets using Ingenuity Pathway Analysis (IPA) analysis

Ingenuity pathway analysis (IPA) evaluates the DEGs and predicts activation or inactivation of pathways. A comparative analysis was done to evaluate the canonical pathways that are activated/inactivated across all subsets in both species using IPA. Pattern Recognition Receptors (PRR) are the first line of defense against any pathogen. The role of RIG-I-like receptors (RLRS) - RIG-1, LGP2 and MDA-5 that sense viral infection, [32] was found to be predominant in CD4^+^, CD8^+^ and CD14^+^ cell subsets at 5 dpv in goats and, in CD4^+^ and CD14^+^ cell subsets of sheep **(Figure 5)**. This RIG-I recognition of viral RNA induces anti-viral state in cells by phosphorylating the IRFs [33] and regulating NF-κB activity through binding to *Nf-κb1* 3’-UTR mRNA [34]. This activation of IRFs by cytosolic pattern recognition receptors was found to be significant in CD4^+^ cells of goats and was triggered, though not significant in CD4^+^ cells of sheep, and in CD8^+^ and CD14^+^ cells of both the species **(Figure 5)**. IRF3 was upregulated in CD4+, CD14+, CD21+ and CD335+ of goats, and in CD335+ of sheep; IRF7 was upregulated in CD4+, CD8+, CD14+ and CD335+ of sheep, and in all cell subsets of goats **(Supplementary File 1)**. IRF7 was also identified to be the most prominent upstream regulator across subsets in both the species. RNA viruses are also recognized by TLR3 (dsRNA) and/or by TLR7/8 (sRNA) [35]. At 5 dpv, role of PRRs in recognition of viruses was found activated in CD14^+^ cells of both sheep and goats, and in CD8^+^ cells of goats. TLR2 and TLR4 were upregulated in CD14^+^ cells of both sheep and goats and in CD8^+^ cells of goats. This TLR signaling results in activation of NF-κB and induction of IFN-inducible genes and co-stimulatory molecules [36].

**Figure 5:**
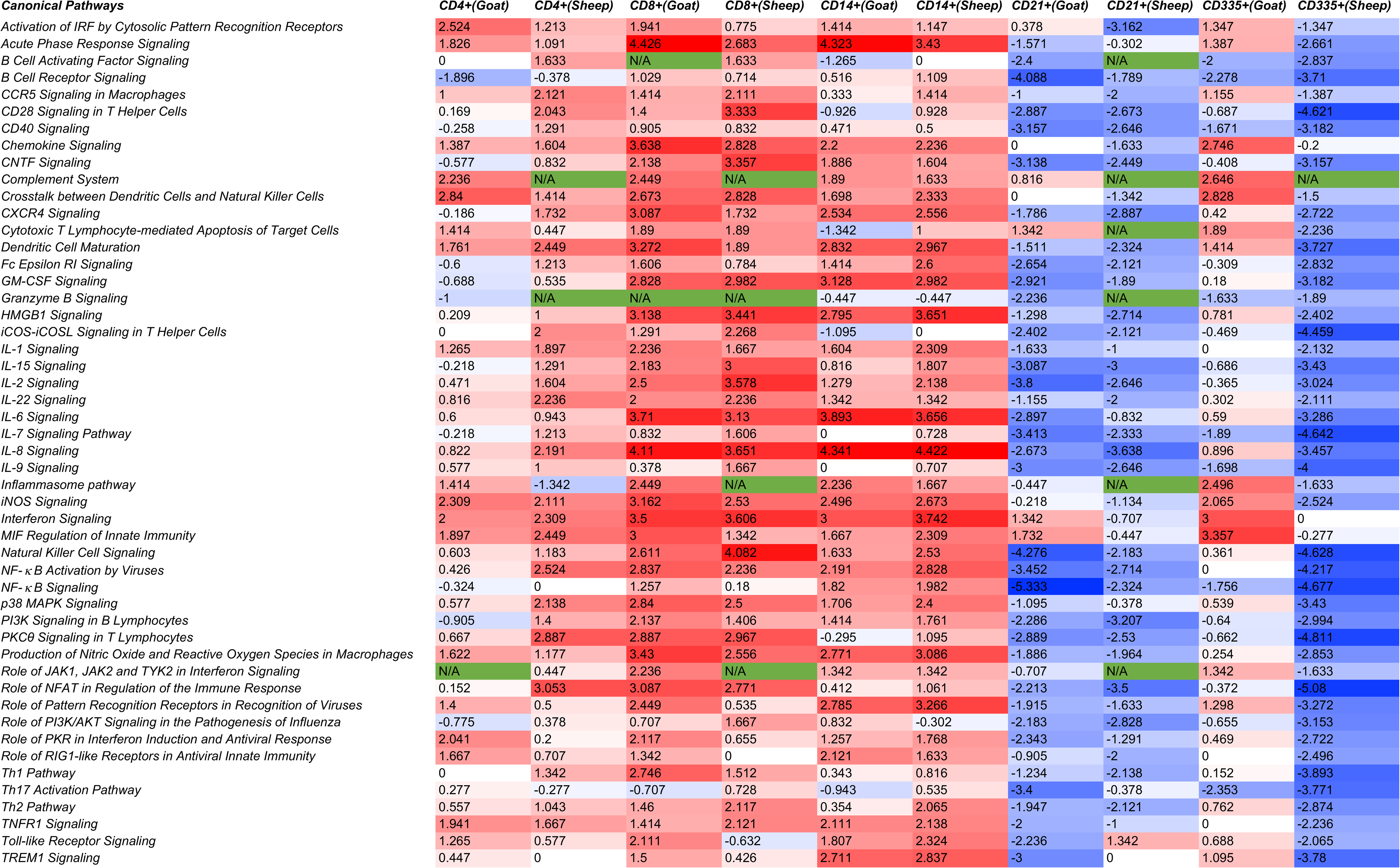
Comparison analysis of canonical pathways related to immunological processes among the subsets of PBMCs in both sheep and goats using IPA. Blue colour represents Z score < 0 and red colour represents Z score > 0. Z score ≥ 2 means activation of canonical pathways and Z score ≤ −2 means inactivation of canonical pathways

The NF-κB activation by viruses was found activated in CD4^+^, CD8^+^ and CD14^+^ cells of sheep and in CD8^+^ and CD14^+^ cells of goats but was inactivated in CD21^+^ cells of both sheep and goats (**Figure 5**). The genes involved in this NF-κB pathway - CD4, LCK, IKK, ERK 1/2, PKR and RIP were upregulated in CD4+ cells of sheep; LCK, RAS, MEKK1, C-RAF, ERK1/2, IκB and CCR5 were upregulated, and CD21 and CXCR5 were downregulated in CD8+ cells of sheep; RAS, PKC, ERK 1/2, IκB, NFκB & PKR were upregulated in CD14+ cells of sheep; CD4, LCK, RAS, PKR, ERK 1/2 and IκB were upregulated and CXCR5 was downregulated in CD8+ cells of goats; RIP, PKR, AKT, IKK, ERK 1/2, IκB & c-RAF were upregulated and CXCR5, CD4 & LCK were downregulated in CD14+ cells of goats(**Supplementary File 1**). NF-κB acts as a mediator of pro-inflammatory and anti - inflammatory gene induction and plays a role in regulating T-cell differentiation and effector function [37]. Several interleukin and chemokine signaling pathways were found activated in CD4^+^, CD8^+^ and CD14^+^ cells of both the species i.e. IL-1 signalling in CD8+ cells of goats and CD14+ cells of sheep; IL-15 signalling in CD8+ cells of sheep and goats; IL-2 signalling in CD8+ cells of sheep and goats and CD14+ cells of sheep; IL-22 signalling in CD4+ cells of sheep and CD8+ cells of sheep and goats; IL-6 signalling in CD8+ and CD14+ cells of sheep and goats; IL-8 signalling in CD4+ cells of sheep, and CD8+ cells and CD14+ cells of sheep and goats, and; chemokine signalling in CD8+ cells and CD14+ cells of sheep and goats **(Figure 5)**.

Dendritic cell (DC) maturation was found significantly activated in CD21+ and CD8+ cells of both the species. DCs are known to present antigenic peptides complexed with MHC class I molecules to CD8-expressing T cells in order to generate cytotoxic cells [38]. The Interferon signalling pathway that is essential for increased cellular resistance to viral infection was found activated at 5 dpv in CD4^+^, CD8^+^ and CD14^+^ cells of both the species. Interestingly, IFN alpha and beta were not dysregulated in any of the subsets in both sheep and goats. The IFN receptors IFNAR1 and IFNAR2 were downregulated in most of the subsets. The absence of expression of type-I interferons in our study suggested IFN-independent ISG stimulation as reported previously for PPR [7]. However, IFNgamma receptors were found to be activated in most of the subsets. Further, most of the canonical pathways were identified to be inactivated in the CD21+ cells. This indicated that the CD21+ cells are activated later for the production of antibodies as significant increase in antibody production against PPRV vaccination was observed 14 dpv [39]. The enrichment (-log p value) of genes in cell mediated immune response and humoral immune response biofunctions was significantly higher in goats than in sheep in CD4^+^, CD14^+^ CD21^+^ and CD335^+^(**Figure 6**).

**Figure 6:**
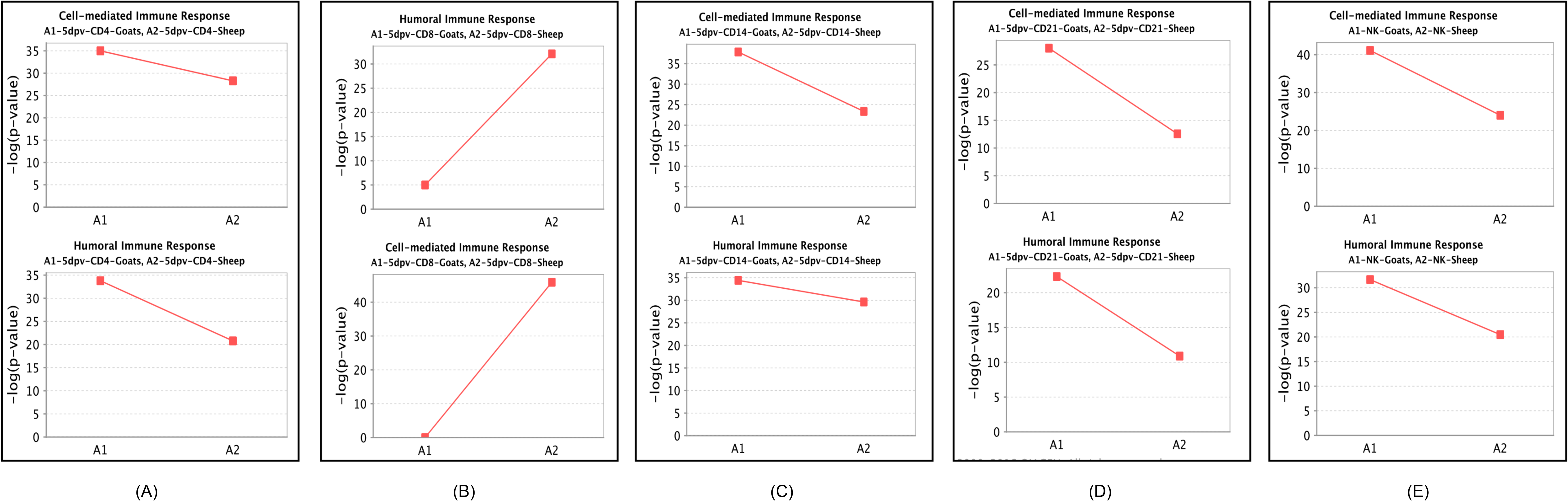
Comparison of significant enrichment (-log p value) of genes in cell mediated immune response and humoral immune response bio functions in (A) CD4+, (B) CD8+, (C) CD14+, (D) CD21+ and (E) NK cells (CD335+) between vaccinated goats and sheep.

### Protein-protein interaction networks

The protein-protein interaction network includes hubs connected with interacting genes. The hubs in a network reflect the functional and structural importance of the network. A total number of 618 and 139 DEGs were found to be commonly expressed in Goats and sheep, respectively in all the subsets (**Figure 2**). On deciphering the interactions between these DEGs and the 8 antiviral molecules (IFIT3, ISG15, MX1, MX2, RSAD2, ISG20, IFIT5 and IFIT1) considered under the knowledge based approach, most of the antiviral molecules formed the hubs in the network. ISG15 in both species was found be the major hub with a connectivity of 75 and 16 in goats and sheep, respectively (**Figure 7A & Figure 7C**). Heatmap of the genes involved in the networks revealed that most of these antiviral genes in both the species are upregulated (**Figure 7B & Figure 7D**).

**Figure 7:**
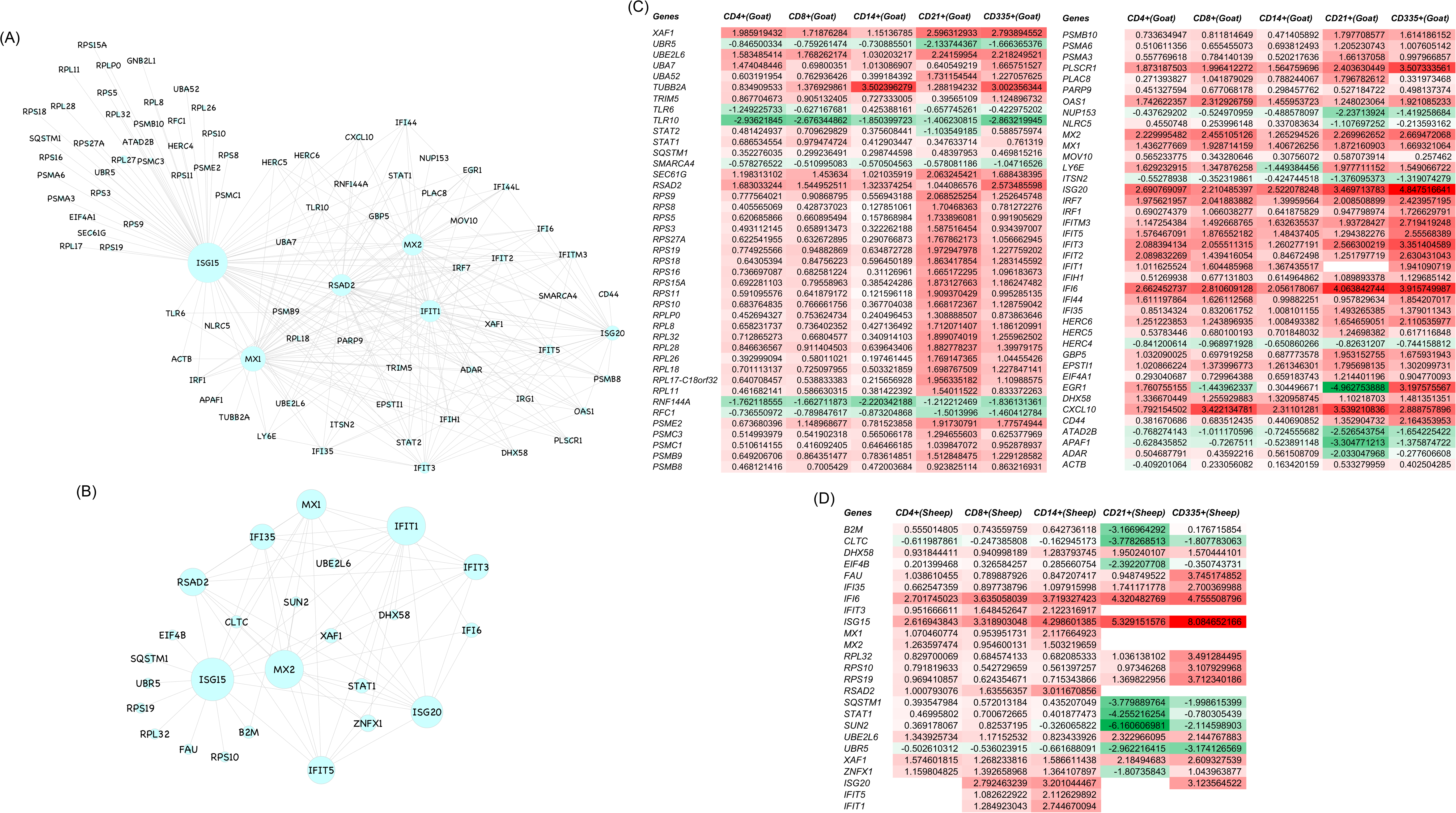
Protein-protein interaction network of antiviral genes - IFIT3, ISG15, MX1, MX2, RSAD2, ISG20, IFIT5 and IFIT1 with the common DEGs across all subsets of each species (A) Goats and (B) Sheep. Size of circle indicates the degree of interaction. (C) Heatmap for fold change (Log_2_FC) values of DEGs involved in the network among the subsets of PBMCs of goats. (D) Heatmap for fold change (Log_2_FC) values of DEGs involved in the network among the subsets of PBMCs of sheep. Green colour indicates downregulation and red colour indicates upregulation.

### Realtime PCR

The key genes identified from RNA-seq data - *DDX58, IFIT3, IRF7, ISG15* and *MX1* were validated by qRT-PCR. The expression of all the validated genes was in concordance with RNA sequencing results **(Table 1)**.

**Table 1.A:** log_2_FC from RNAseq and qRT-PCR of PBMC subsets isolated from sungri/96 vaccinated Goat

|  | CD4 |  | CD8 |  | CD14 |  | CD21 |  | CD335 |  |
| --- | --- | --- | --- | --- | --- | --- | --- | --- | --- | --- |
| Gene | log<sub>2</sub>FC from qRT-PCR | log<sub>2</sub>FC from RNAseq | log<sub>2</sub>FC from qRT-PCR | log<sub>2</sub>FC from RNAseq | log<sub>2</sub>FC from qRT-PCR | log<sub>2</sub>FC from RNAseq | log<sub>2</sub>FC from qRT-PCR | log<sub>2</sub>FC from RNAseq | log<sub>2</sub>FC from qRT-PCR | log<sub>2</sub>FC from RNAseq |
| DDX58 | 1.179333333 | 1.163823481 | 1.123333333 | 1.126889098 | 0.980666667 | 0.897711159 |  |  | 0.97 | 0.866777509 |
| IFIT3 | 2.308666667 | 2.088394134 | 2.12 | 2.055511315 | 0.869802704 | 1.260277191 | 2.630251215 | 2.566300219 | 3.071333333 | 3.351404589 |
| IRF7 | 2.005631332 | 1.975621957 | 1.945542755 | 2.041883882 | 1.051373011 | 1.39959564 | 1.936792165 | 2.008508899 | 2.627666667 | 2.423957195 |
| ISG15 | 3.193989334 | 3.483488156 | 3.214456762 | 3.642171919 | 1.508623454 | 2.749599382 | 5.020430272 | 4.958535352 | 4.033333333 | 4.817152957 |
| MX1 | 1.164756699 | 1.436277669 | 1.825847584 | 1.928714159 | 1.289272029 | 1.406726256 | 2.063278147 | 1.872160903 | 1.556666667 | 1.669321064 |

**Table 1.B:** log_2_FC from RNAseq and qRT-PCR of PBMC subsets isolated from sungri/96 vaccinated Sheep

|  | CD4 |  | CD8 |  | CD14 |  | CD21 |  | CD335 |  |
| --- | --- | --- | --- | --- | --- | --- | --- | --- | --- | --- |
| Gene | log<sub>2</sub>FC from qRT-PCR | log<sub>2</sub>FC from RNAseq | log<sub>2</sub>FC from qRT-PCR | log<sub>2</sub>FC from RNAseq | log<sub>2</sub>FC from qRT-PCR | log<sub>2</sub>FC from RNAseq | log<sub>2</sub>FC from qRT-PCR | log<sub>2</sub>FC from RNAseq | log<sub>2</sub>FC from qRT-PCR | log<sub>2</sub>FC from RNAseq |
| DDX58 | 1.311 | 1.443982294 | 2.006666667 | 1.880321932 | 1.806666667 | 1.630166862 |  |  |  |  |
| IFIT3 | 0.536779277 | 0.951666611 | 1.696666667 | 1.648452647 | 1.923333333 | 2.122316917 |  |  |  |  |
| IRF7 | 2.214341863 | 2.174678699 | 1.843333333 | 2.320513832 | 3.233333333 | 3.017430161 |  |  | 3.153333333 | 3.669000639 |
| ISG15 | 2.781512248 | 2.616943843 | 2.796666667 | 3.318903048 | 3.126666667 | 4.298601385 | 4.142500178 | 5.329151576 | 5.643333333 | 8.084652166 |
| MX1 | 1.34357254 | 1.070460774 | 1.01 | 0.953951731 | 2.186666667 | 2.117664923 |  |  |  |  |

## Discussion

Vaccines protect against an infectious agent by inducing cells or molecules capable of rapidly controlling their replication or by inactivating their toxins. Primarily, vaccines trigger an inflammatory reaction, mediated by cells of the innate immune system - dendritic cells, monocytes and neutrophils. These cells recognize PAMPs through PRRs to get activated to produce cytokines and chemokines [40], [41], [42], [43]. This inflammatory microenvironment is essential for differentiation of monocytes to macrophages, and for activation and migration of dendritic cells into the draining lymph nodes [44]. In the absence of this inflammatory response the dendritic cells remain immature and the naive T cells in the lymph nodes do not differentiate into CD4+ T cells. It is evident that PPRV - Sungri/96 live attenuated vaccine triggers activation of the innate immune system after it is phagocytosed by monocytes/dendritic cells at the site of administration [45]. This RNA virus may be then recognized by TLR3/7 on the endosome or by the RIG-1 or MDA5 in the cytosol to induce an inflammatory response. This induction of inflammatory response is evident in both sheep and goats with the triggering of several pathways viz. role of RIG1-like receptors in antiviral innate immunity, role of pattern recognition receptors in recognition of viruses, production of nitric oxide and reactive oxygen species in macrophages, NF-KB activation by viruses, and several IL signaling pathways in CD14+ cells. This triggering in inflammatory response is much needed for the activation of dendritic cells and monocytes, and for further draining of these cells to the nearest lymph node where naive T cells are activated [44]. This activation of T cells is clearly seen by the activation of pathways in CD8+ (T-cytotoxic) and CD4+ (T-helper) cells.

Out of the several pathways activated in both CD4+ and CD8+ cells, NOD-like receptor signaling pathway, Th1 and Th2 cell differentiation, T cell receptor signaling pathway and Th17 cell differentiation were found significantly enriched in both the species. The differentiation of T cells to Th1 and Th2 is crucial for inducing the immune response. Th1 cells stimulate cellular immune response, participate in the inhibition of macrophage activation and stimulate B cells to produce IgM and IgG1 [46]. Th2 stimulates humoral immune response, promotes B cell proliferation and induces antibody production [46]. The distinct subsets of helper T cells - TH1, TH2 and TH17, are effective at protecting against pathogens [47]. Additionally, activation of C-type lectin receptors (CLRs) signaling in CD8+ cells of Sungri/96 vaccinated sheep and goats and in CD4+ cells of goats indicates induction of adaptive immune response. C-type lectin receptors (CLRs) are important pattern recognition receptors involved in recognition and induction of adaptive immunity to viruses [48]. However, in CD21+ cells most of the pathways were found inactivated/not activated as 5 dpv may be too early a time point to detect activation in the CD21+ cells.

NK cells (CD335+) are known to mediate both innate immune and adaptive immune responses by modulating both CD8+ and antibody production [49]. In this study most of the pathways - interferon signaling, crosstalk between dendritic cells and natural Killer cells, chemokine signaling, inflammasome pathway, iNOS signalling and complement system in NK cells were found activated in vaccinated goats than in sheep. Upregulation of RIG-1 and MDA5 in NK cell of goats reflects setting off of the innate immune response [50]. Also, activation of interferon signaling pathway in infected NK cells of goats suggests evoking of both the innate and adaptive immune responses [51]. The activation of iNOS signaling invokes immune response in virus infected cells [52].The activation of complement system in NK cells aids in antibody production by bridging both innate and adaptive immune response [53]. Upregulation of CD69, NKp30, FAS & TNFR2 and activation of crosstalk between dendritic cells and natural killer cells pathway, in goats must be embarking innate immune response, followed by adaptive response on antigen presentation after vaccination. The activation and triggering of several pathways in NK cells of goats at this early time point may be because of Sungri/96 vaccine strain being of goats origin and that the activation of these pathways at a later time point in sheep cannot be ruled out.

The network of antiviral molecules (IFIT3, ISG15, MX1, MX2, RSAD2, ISG20, IFIT5 and IFIT1) with the DEGs commonly expressed in the subsets in both sheep and goats, reflected ISG15 as a major hub. The network was found to be dense in goats in comparison to sheep. ISG15 is one of the most highly induced ISGs in viral infections [25], [54] and was also found to be directly induced by IRF3/IRF7, independent of IFNs [55], [56], [57]. It is an ubiquitin-like protein that covalently attaches to target proteins in a process known as ISGylation [25], [58]. HERC5 is considered as the major ligating enzyme in ISGylation. This ISGylation of viral proteins was reported to have an inhibitory effect on the viral infection [59], whereas ISGylation of host proteins leads to either activation [59] or increase in stability [60]. HERC6 instead of HERC5 is considered as the major ligating enzyme in mice [61]. In our study, HERC5 and HERC6 were found upregulated in goats. Further, the antiviral gatekeeper MX1 acts prior to genome replication at an early post entry step of the virus life cycle. Similarly, MX2 specifically targets viral capsid and affects nuclear entry of the HIV-1 [25], [62], [63], [64]. IFIT family (IFN-induced protein with tetratricopeptide repeats) are a group of ISGs that inhibit virus replication by binding and regulating the functions of cellular and viral proteins and RNAs [65]. IFITs were also characterized to play a critical role in protecting hosts from viral pathogenesis. RSAD2, also known as Viperin, is the another most highly induced antiviral effector found in ER and ER-derived lipid droplets [66]. RSAD2 was characterized to have various modes of antiviral action to inhibit enveloped viruses [67]. It can also affect virus life cycle at an early stage by inhibiting RNA replication [68]. All these genes - MX1,MX2,IFIT1, RSAD2, IFIT3 and IFIT5 were found upregulated in both sheep and goats suggesting a strong antiviral response in both the species.

It is important to note the in our study both sheep and goats survived PPRV virulent virus challenge post vaccination, indicating an adequate immune response to counter the virus. In an independent study it was reported that Sungri/96 vaccine is equally potent in both sheep and goats [69]. In our study, though there are a few differences in the systems biology across cells (specially the NK cells) between sheep and goats, the co-ordinated response that is inclusive of all the cell subsets was found to be towards induction of strong innate immune response, which is needed for an appropriate adaptive immune response.

## Author contributions

RKS, BPM, and RKG conceived and designed the research. KKR and DM performed the vaccine testing experiment. SAW and ARS conducted the wet lab work. SAW, MRP, RINK and RKG analyzed the data. SAW, MRP, RKG, APS, and BM helped in manuscript drafting and editing. RKS, BPM, and RKG proofread the manuscript.

## Acknowledgment

This study was supported in part by Centre for Agricultural Bioinformatics (ICAR-IASRI) (CABin/100644/16103/ 801/10133) and SubDIC (BTISnet), ICAR-IVRI. We also thank Department of Biotechnology, Ministry of Science and Technology (DBT) for providing fellowship and contingency for students – Sajad Ahmad Wani (DBT/2014/IVRI/171), Manas Ranjan Praharaj (CSIR:09/1150(0015)/2019-EMR-I) and Amit Ranjan Sahu (DBT/2014/IVRI/170).

---

**Supplementary Figure 1:** Purity of PBMCs subsets - T helper cells (CD4+), T cytotoxic cells (CD8+), monocytes (CD14+), B lymphocytes (CD21+), and natural killer cells (CD335+) by flow cytometry. ‘PBMCs’ means unstained cells. ‘PBMC’ with Ab’ means before magnetic beads cell separation. Enriched cells mean post magnetic beads cell separation.

**Supplementary Figure 2:** Heatmap for fold change (Log_2_FC) value of common immune DEGs between goats and sheep for each subset of PBMCs: (A) CD4+, (B) CD8+, (C) CD14+, (D) CD21+ and (E) CD335+. Green colour indicates downregulation and red colour indicates upregulation.

**Supplementary Figure 3:** Functional annotation related to immunological processes of genes expressed in all the subsets or in one subset of PBMCs associated with innate immune response (A) Goats (B) Sheep

**Supplementary File 1:** Individual list of DEGs expressed in the subsets of goats and sheep

**Supplementary File 2:** List of 5512 and 5297 DEGs expressed across the subsets in goats and sheep, respectively

